# Specimen alignment with limited point-based homology: 3D morphometrics of disparate bivalve shells (Mollusca: Bivalvia)

**DOI:** 10.1101/2022.03.04.482893

**Authors:** Stewart M. Edie, Katie S. Collins, David Jablonski

## Abstract

Comparative morphology fundamentally relies on the orientation and alignment of specimens. In the era of geometric morphometrics, point-based homologies are commonly deployed to register specimens and their landmarks in a shared coordinate system. However, the number of point-based homologies commonly diminishes with increasing phylogenetic breadth. These situations invite alternative, often conflicting, approaches to alignment. The bivalve shell (Mollusca: Bivalvia) exemplifies a homologous structure with few universally homologous points—only one can be identified across the Class, the shell ‘beak.’ Here, we develop an axis-based framework, grounded in the homology of shell features, to orient shells for landmark-based, comparative morphology. As the choice of homologous points for alignment can affect shape differences among specimens, so can the choice of orientation axes. Analysis of forty-five possible alignment schemes finds general conformity among the shape differences of ‘typical’ equilateral shells, but the shape differences among atypical shells can change considerably, particularly those with distinctive modes of growth. Each alignment implies a hypothesis about the ecological, developmental, or evolutionary basis of morphological differences, but we recognize one alignment in particular as a continuation of the historical approaches to morphometrics of shell form: orientation via the hinge line. Beyond bivalves, this axis-based approach to aligning specimens facilitates the comparison of continuous differences in shape among many other phylogenetically broad and morphologically disparate samples.

## 2. Introduction

Comparative morphology depends on how organisms are oriented, or aligned. For a simplistic example, a kiwi’s beak is relatively long for a bird when measured from the tip to the base of the skull, but rather short when measured from the tip to the nostrils (an alternative definition of beak length; Borras et al. 2000). Thus, the choice of anatomical reference points can profoundly alter our interpretations of evolutionary morphology. Alignments commonly use point-based aspects of homologous features—the junction of the kiwi’s beak with the cranium (a Type I landmark; Bookstein 1992) and the distal-most point of the beak, the tip (a Type III landmark). Closely related organisms tend to have more of these homologous points, allowing for a straightforward alignment and comparison of their shapes. Alignment on strict, point-based homology becomes more problematic with increasing phylogenetic distance, however, as the number of homologous features invariably diminishes (Bardua et al. 2019).

Bivalve mollusks have become a model system for macroevolution and macroecology (Jablonski et al. 2017; Edie et al. 2018; Crame 2020), but their strikingly disparate body plans complicate Class-wide morphologic comparisons using strict homology (Cox et al. 1969). Inimical to triangulation and thus alignment via landmarks, the valve of the bivalve shell—the most widely available feature of the animal today and through the fossil record—has only one homologous *point*; the apex of the beak, which is the origin of growth of the embryonic shell at the apex of the beak (Carter et al. 2012:21; Figure 1). Homology-free approaches can be useful for comparing the shapes of shell valves when anatomical orientation is either unknown or uncertain (Bailey 2009); but wholesale substitution of shape, i.e. analogy, for homology complicates the evolutionary interpretation of morphological differences. Despite the lack of multiple homologous *points* on the shell valve across the Class, a number of its *features* are homologous and can facilitate comparisons. Here, we apply principles of bivalve comparative morphology to develop a framework for aligning shell valves (hereafter ‘shells’) across the Class, thus enabling phylogenetically extensive analyses of their shapes using geometric morphometrics despite the remarkable range of body plans across the clade.

**Figure 1.**
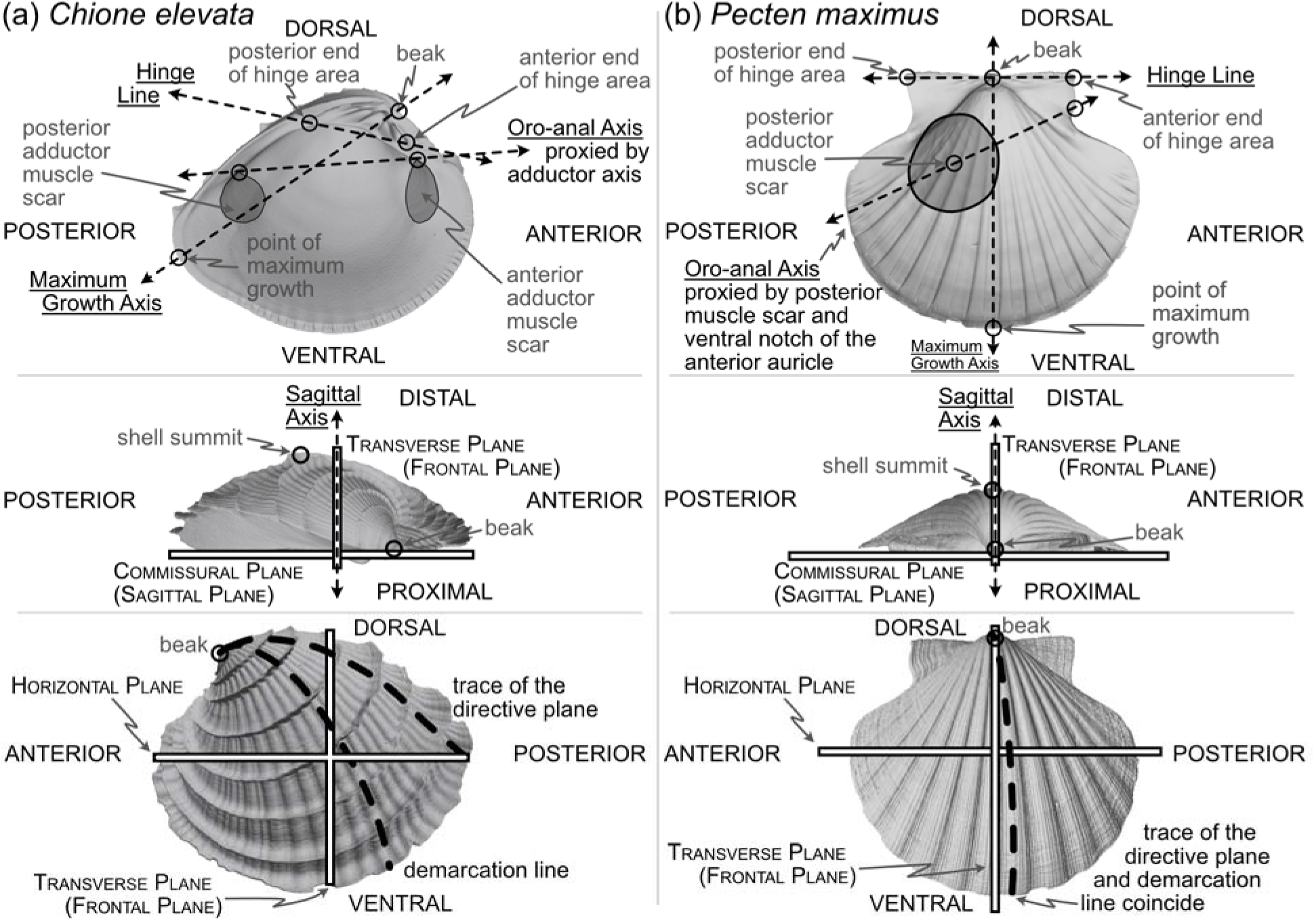
Positions of shell features, axes, and planes as mentioned and defined in text for: (a) a helicospiral shell *Chione elevata* (Say 1822), and (b) a more planispiral shell *Pecten maximus* (Linnaeus 1767).

### 2.1 Approaches to orienting the bivalve shell

Many body directions, axes, lines, and planes have been defined for bivalves (see Cox et al. 1969; Bailey 2009; Carter et al. 2012)—some related to features of the shell (an accretionary exoskeleton composed of calcium carbonate; Marin et al. 2012), and others to features of the soft body (the digestive tract, foot, byssus, muscles, etc.). Separation into these ‘shell’ and ‘body’ terms is a false (Stasek 1963*a*) but convenient dichotomy (Yonge 1954): the shell is generated by, and remains attached to, the soft body, but their morphologies can become decoupled (Yonge 1954; Edie et al. 2022). Still, both shell and body features are required for orientation via homology (Stasek 1963*a*; cf. Bailey 2009). Given our goal of aligning the shell for geometric morphometrics across broad phylogenetic scales and through the fossil record, we focus on orientations that can be inferred from this part alone, but we must use a critical body axis to fully determine orientation, the anteroposterior axis. Thus, shell features used for alignment can be divided into two classes: (1) intrinsic characteristics of the shell relating to its geometry, growth, and biomechanics, and (2) proxies of the body recording the positions of the soft anatomy including the adductor muscle scars, pallial line, byssal notch, pedal gape, siphon canal, and more.

#### 2.1.1 Orientation via intrinsic characteristics of the shell

Although the only homologous point on the shell across all bivalves is the apex of the beak (hereafter ‘beak’, Figure 1), aspects of the shell’s biomechanics, such as its hinge axis, and its accretionary growth permit orientation via homology. The planes and lines proposed to describe shell geometry and growth have been criticized for their lack of homology (Stasek 1963b:226) and ubiquity across the class (Lison 1949:62; Owen 1952:149; Carter 1967:272). We discuss two such features here, the directive plane and demarcation line (Figure 1 and defined below), but we also address and test two alternative means of orienting shells that combine the shell’s geometry and growth: the maximum growth axis and the shape of the shell commissure.

Related to the shell’s biomechanics, the hinge has been treated as a “fixed dorsal region” (Yonge 1954:448; see also Jackson 1890:282), later redefined to reflect the position of the mantle isthmus bridging between the two valves as a universally dorsal-directed feature (Cox et al. 1969:79). Beyond its determination of dorsoventral directionality, the hinge, specifically the hinge axis defined as the “ideal line drawn through the hinge area, and coinciding with the axis of motion of the valves” (Jackson 1890:309), is a Class-wide feature that can constrain one of the three Cartesian axes required for alignment. In a strictly mechanical sense, the ligament, and not the hinge teeth, directs the orientation of the axis of motion (Trueman 1964:56; Cox et al. 1969:47; Stanley 1970:47). However, the hinge area, which includes the teeth, is hypothesized to be analogous in function (Cox et al. 1969:47)—guiding the two valves into alignment during closure—and homologous in its origin (Scarlato and Starobogatov 1978; Waller 1998; Fang and Sanchez 2012). Thus, our definition of the **hinge line** indicates the longest dimension of the hinge area (see ‘hinge’ in Carter et al. 2012:74 and Figure 1); for quantitatively aligning shells, the hinge line is determined by the two farthest apart articulating elements of the hinge area, excluding lateral teeth, which are variably present among heterodont species (e.g. Mikkelsen et al. 2006:493; Taylor and Glover 2021); definition proposed here is a synthesis of the discussions in Cox et al. 1969:81 and Bradshaw and Bradshaw 1971). Thus, by directing the orientation of the horizontal plane which divides the body into dorsal (towards the beak) and ventral (towards the free edge of the shell) territories, the hinge line can also proxy the anteroposterior axis (Figure 1, Cox et al. 1969:81 and discussion below).

Related to the geometry and growth of the shell, the **directive plane** (Lison 1949) was proposed as the only plane passing through the shell that contains the logarithmic planispiral line connecting the beak to a point on the ventral margin (Figure 1); all other radial lines would be logarithmic turbinate spirals (or ‘helicospirals’; Stasek 1963b:217). In other words, on a radially ribbed shell, there may be a single rib that lies entirely on a plane when viewed from its origin at the umbo to its terminus on the ventral margin; that plane is orthogonal to the commissural plane for planispiral shells (e.g. many Pectinidae, Figure 1), but lies at increasingly acute angles to the commissural plane with increasing tangential components of growth (i.e. geometric torsion; see the trace of the directive plane on *Chione* in Figure 1 and examples in Cox et al. 1969:86-Figs. 70-71). In theory, the directive plane could be used to orient the dorsoventral axis of the shell, but in practice, the feature is not universal across shell morphologies (e.g. the strongly coiled *Glossus humanus* [as *Isocardia cor]* in Lison 1949:62; Owen 1952:149). Cox et al. (1969:87) also remark that the plane cannot “be demonstrated easily by visual inspection if the shell lacks radial ribbing, except in rare specimens with an umbonal ridge that proves to lie within the directive plane.” Difficulty in application is no excuse to avoid an approach, but the non-universality of this feature renders it inapplicable to Class-wide comparisons of shell shape.

Owen (1952) proposed an alternative to the directive plane: the **demarcation line** (Figure 1; originally termed the ‘normal axis’ but re-named by Yonge 1955:404). As with the directive plane, the demarcation line serves to orient the dorsoventral direction and separate the shell into anterior and posterior ‘territories’ (Yonge 1955:404; Morton and Yonge 1964:40), but its definition has been variably characterized in geometric and/or anatomical terms. Per Owen (1952:148), the demarcation line can “be considered with reference to three points: the umbo, the normal zone of the mantle edge and the point at which the greatest transverse diameter of the shell intersects the surface of the valves.” Yonge (1955:404), acknowledging correspondence with Owen, described the demarcation line as: “the projection onto the sagittal plane of the line of maximum inflation of each valve… starting at the umbones.… i.e. the region where the ratio of the transverse to radial component in the growth of the mantle/shell is greatest.” Carter et al. (2012:52) provided perhaps the clearest description as the line defining the “dorsoventral profile when the shell is viewed from the anterior or posterior end.” However, Stasek (1963*b*) demonstrated the difficulty in measuring this line; note the nearly orthogonal orientations of the empirically determined demarcation line on *Ensis* (Stasek 1963*b*:225-Fig.5a) compared to its initially proposed position (Owen 1952:148-Fig. 5). Stasek’s empirical approach, coupled with the revised definition of Carter et al., is tractable with today’s 3D-morphology toolkit. But, critically, this definition depends on the direction of the anteroposterior axis, which itself is variably defined (see discussion in next section). Thus, definitionally driven shifts in the direction of the anteroposterior axis can alter the trace of the demarcation line. Owen’s initial definition is independent of the anteroposterior axis, but as Stasek demonstrated, its identification can be unreliable. Thus, high degrees of digitization error for the demarcation line may confound comparisons of shell shape, and we do not include the demarcation line as a feature for aligning shells across the Class.

Both the directive plane and the demarcation line attempt to orient the shell on aspects of its geometry that are intrinsic to its growth. A similar and more reliably determined approach may be orientation to the **maximum growth axis** (i.e. **line of greatest marginal increment** sensu Owen 1952; Figure 1). The maximum growth axis is the straight line that connects the origin and terminus of the trace of maximum growth across the shell surface. This trace connects the beak to the ventral margin along a perpendicular path to the most widely spaced commarginal growth increments (as such, this definition appears to have similar properties to the trace of the directive plane on the shell surface). But, as for the directive plane and the demarcation line, the maximum growth axis can be prone to measurement error without fitting a formal model of shell growth (e.g. those of Savazzi 1987; Ubukata 2003), and should therefore be used with caution. However, a reasonable and reliably measured proxy for this axis is the line lying on the commissural plane that originates at the beak and terminates at the furthest point on the shell commissure. Thus, this axis can indicate the dorsoventral orientation of the shell.

Orientation using the **shape of the shell commissure** offers, arguably, the most reliably determined approach that uses intrinsic properties of the shell (Figure 2a). Given the accretionary growth of the shell, points on the commissure—the homologous leading edge of shell growth (Vermeij 2013)—are geometrically homologous, or correspondent (Bookstein 1991; Gunz et al. 2005). Valve handedness is still required to ensure that compared valves are from the same side of the body (i.e. left vs. right), which requires anteroposterior directionality (see below). This alignment thus orients shells using geometric correspondence based on homology of growth.

**Figure 2.**
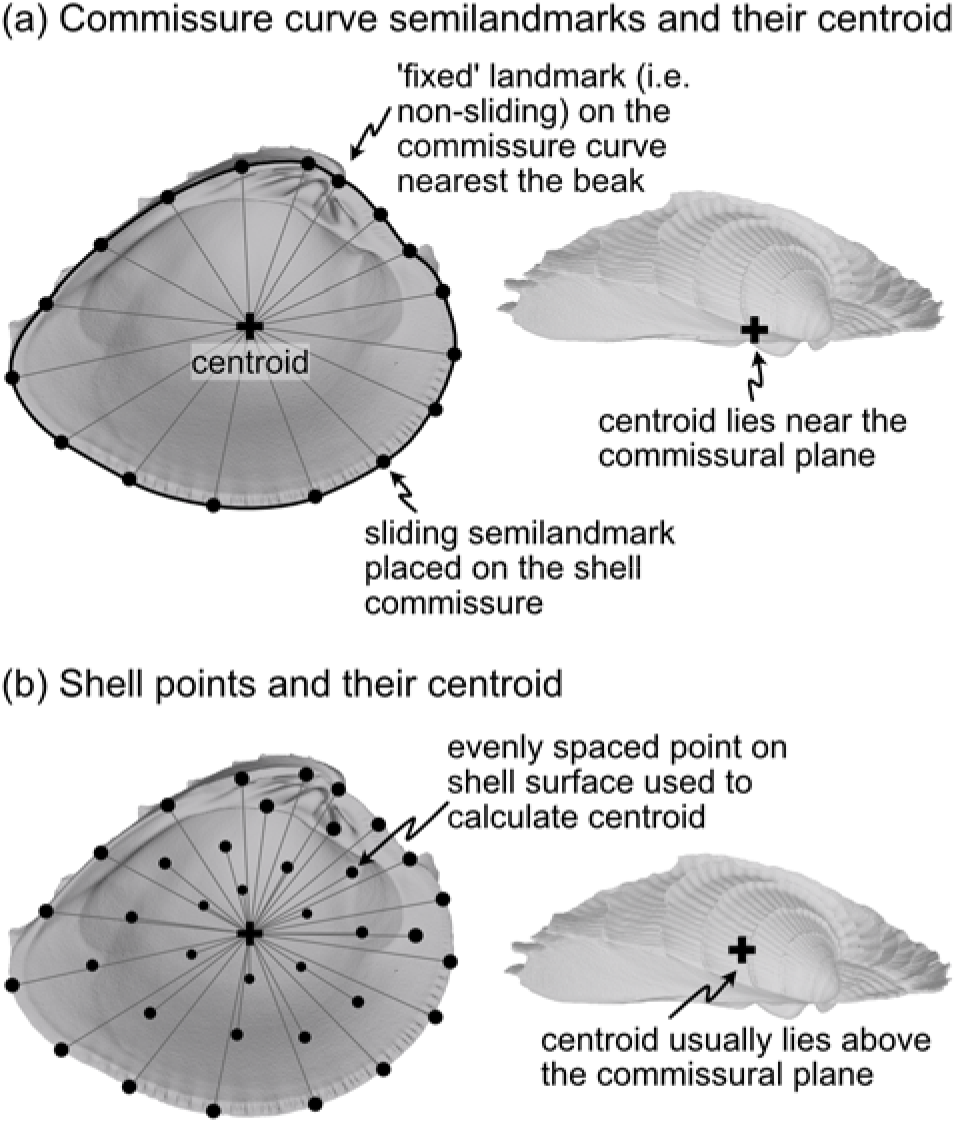
(a) Representation of shell commissure curve, its centroid, and the semilandmarks used in the COMM orientation scheme. Analyses use 50 sliding semilandmarks on the commissure curve, but only a subset is shown here for clarity. (b) Equally spaced points on the shell surface placed using a Poisson Disc sampler (*Rvcg::vcgSample*, Schlager 2017) and their centroid. The number and location of vertices on triangular meshes can vary, which strongly influences the calculation of centroid size. Analyses use 2000 equally spaced points to minimize this issue (only a subset of those points shown here). Figured shell is *Chione elevata*.

The sagittal axis is crucial to the shell’s three-dimensional alignment and is likely the least controversially defined. This axis is the pole (=normal) to the sagittal plane, which lies parallel to the commissural plane defined as: “the more proximal part of the line or area of contact of the two shell valves” (Carter et al. 2012:38). Therefore, the sagittal axis is parallel to the frontal and horizontal planes (Figure 1). The proximal direction is towards the commissural plane and the distal direction is towards the shell’s summit: the point on the shell that is maximally distant from the commissural plane (Figure 1, Cox et al. 1969:108; Carter et al. 2012:177). If valve handedness (i.e. left vs. right laterality) and the directionality of the dorsoventral and anteroposterior axes are known, then this axis is rarely required for orientation. However, certain definitions of the anteroposterior and dorsoventral axes are not constrained to be orthogonal (e.g. in monomyarian taxa with the anteroposterior axis defined as the oro-anal axis, see below, and the dorsal ventral axis as the axis of maximum growth—*Pecten* in Figure 1); as the axes representing the anteroposterior and dorsoventral directions become more parallel, then the sagittal axis becomes an increasingly important safeguard against the inversion of the proximal-distal direction in quantitative alignments.

#### 2.1.2 Orientation via the soft-body as reflected on the shell

Anteroposterior directionality is the third Cartesian axis required for orienting the bivalve shell in three-dimensions. The positions of the mouth (anterior) and anus (posterior) ultimately determine the anteroposterior axis (Jackson 1890, ‘preferably’ described as the ‘oro-anal’ axis in Cox et al. 1969:79), but the exact positions of these two soft-body features are rarely recorded directly on the shell. Thus, tactics for determining the polarity, if not the precise bearing, of the anteroposterior axis have relied on lineage or body-plan specific proxies—shell features that are assumed to correlate with positions of the soft-body anatomy (e.g. positions of the adductor muscle scars, Figure 1a, Cox et al. 1969:79). Disparate body plans necessitate taxon-specific rules for orientation, such that determining the anterior and posterior ends of the shell requires a mosaic approach. For example, there are at least three definitions of the anteroposterior axis in dimyarians alone (Bailey 2009:493), which necessarily differ from those of monomyarians considering the reliance on two, instead of one, adductor muscle scars. For those monomyarians, which commonly have lost the anterior adductor (Yonge 1954; but see loss of the posterior adductor in the protobranch Nucinellidae, Allen and Sanders 1969; Glover and Taylor 2013), additional shell features are used to orient the anteroposterior axis. In pectinids, the byssal notch of the anterior auricle proxies the location of the mouth (Figure 1b), but in ostreids, the mouth is more centrally located under the umbo, near the beak (Yonge 1954:448).

Lineage or body-plan specific definitions help with anteroposterior orientation of shells that lack point-based homology (e.g. two muscle scars vs. one), but they still rely on proxies for the position of soft-body features that may not be determined for taxa known only from their shells, e.g. some fossils (Bailey 2009). Hypothesizing anteroposterior orientation using phylogenetic proximity to extant clades may help, but this approach should be used with caution in given the lack of direct anatomical evidence—especially when phylogenetic affinities are either unknown or distant, as for many Paleozoic taxa (Bailey 2009). Nor is it advisable to assume the precise bearing of the anteroposterior axis is identical to another, well-defined axis such as the hinge line (see variable bearings of the anteroposterior axis and hinge *axis* in Cox et al. 1969:80-Fig. 64). However, if the phylogenetic or temporal scope of an analysis precludes the determination of the anteroposterior axis using homologous body features with geometric correspondences (e.g. inclusion of dimyarian and monomyarian taxa), then the hinge line offers a universal proxy; that way, multiple features can be used to determine the anterior and posterior ends of the shell (Cox et al. 1969).

### 2.2 Alignment of shells for geometric morphometrics

Our challenge here is to reconcile the many means of orienting bivalve shells discussed above into alignment schema for geometric morphometrics. We compare the differences in shell shape produced by the five orientations listed below. All five orientations use the sagittal axis (SX) to determine the lateral orientation of the shell. Precise definitions of landmark placement for each axis are provided in the Methods.

- **SX-HL-oHL.** Anteroposterior orientation determined by the hinge line (HL); dorsoventral orientation determined by the orthogonal line to the HL within the commissural plane (oHL). This alignment emulates the orientation scheme for measuring shell height, length, and width—the most common and widely applicable framework for comparing shell morphology (Cox et al. 1969:81–82; Kosnik et al. 2006).
- **SX-OAX-oOAX.** Anteroposterior orientation determined by the proxied positions of the mouth and anus using shell features (oro-anal axis, OAX); dorsoventral orientation determined by the orthogonal line to the OAX within the commissural plane (oOAX). Similar to SX-HL-oHL, this alignment largely determines orientation by a single axis, the OAX, which has also been used to frame linear measurements of shell morphology (e.g. Stanley 1970:19).
- **SX-HL-GX.** Anteroposterior orientation determined by the hinge line (HL); dorsoventral orientation determined by the maximum growth axis (GX). This alignment allows an aspect of shell growth to affect its orientation and thus the degrees of morphological similarity among specimens.
- **SX-HL-GX-OAX.** Anteroposterior orientation determined by the directions of both the HL and OAX; dorsoventral orientation determined by GX. This ‘full’ alignment scheme allows axes derived from intrinsic characteristics of the shell and the body to affect orientation.
- **SX-COMM.** Anteroposterior and dorsoventral orientation determined by the shape of the commissure curve, with the initial point nearest the beak (Figure 2a). This alignment uses the geometric correspondence of semilandmarks on the commissure that capture the relationship between its shape and growth.

Procrustes superimposition (or Procrustes Analysis) is the workhorse of geometric morphometrics—aligning shapes by translation to a common origin, scaling to common size, and rotation to minimize relative distances of landmarks (see variants thereof in Zelditch et al. 2012). While we are most concerned with assessing the effects of rotation using the five alignments described immediately above, choices of translation and scaling can also influence shape differences. Thus, we consider all combinations of parameter values for each step in the Procrustes Analysis. As there is arguably no objective criterion to determine which alignment best suits bivalve shells, we discuss the benefits and drawbacks of each approach and quantitatively compare the similarities of resulting alignments. Ultimately, we use this exercise to propose a best practice for aligning bivalve shells and comparing their shapes—a process that may be of use for workers in other, similarly disparate morphological systems that lack high degrees of point-based homology.

## 3. Methods

### 3.1 Dataset

We adopt the style of previous approaches to studying bivalve orientation and use a dataset of morphological end-members to illustrate the effects of different alignment schemes (e.g. Owen 1952; Yonge 1954; Stasek 1963*a*). Eleven species that represent most major body plans were selected to proxy the morphological and anatomical disparity across the evolutionary history of the Class (Table S1). Bivalves with highly reduced shells or those that form part of a larger structure (tubes and crypts) are not directly analyzed here, but we consider their fit to the alignments in the Discussion.

One valve from an adult individual of each species was sampled from museum collections (see Acknowledgments). Nine of eleven individuals are equivalve, and because we do not examine details of dentition, their left and right valves are operationally mirror images of each other. The inequivalve taxa included here (*Pecten, Ostrea*) primarily differ in terms of inflation (height above the commissural plane); for the purposes of our analysis, the location of key features such as the hinge area and adductor muscle scars are similar enough that using either valve gives a similar orientation. Valves were scanned using micro-CT at the University of Chicago’s Paleo-CT facility, and three-dimensional, isosurface, triangular-mesh models were created in VG Studio Max and cleaned in Rvcg (Schlager 2017) and Meshmixer. Landmarks were placed using ‘Pick Points’ in Meshlab (Visual Computing Lab ISTI – CNR 2019). Meshes, landmarks, and code necessary to reproduce the analyses here are provided in the Supplemental Material.

### 3.2 Aligning bivalve shells in a geometric morphometrics framework

#### 3.2.1 Scaling

Procrustes Analysis scales objects to a common size, and three alternative scalings are considered here: (1) the **centroid size of the shell** (Figure 2b), (2) the **centroid size of the commissure** (Figure 2a), and (3) the **volume of the shell**. The centroid size of the shell reflects the 3D footprint of the shell, and the centroid size of the commissure reflects the size of the shell’s growth front. Shell volume—the amount of calcium carbonate—may be less correlated with shell shape than the other two measures, and may therefore reveal shape differences not intrinsically linked to size.

#### 3.2.2 Translation

After scaling, Procrustes Analysis translates objects to a common origin. Objects are typically ‘centered’ by subtracting the centroid of the landmark set (mean X, Y, and Z coordinate values per object) from each landmark coordinate, thus shifting the center of each landmark set to the origin (X=0, Y=0, Z=0). Three points are considered for translation here: (1) the **beak** (Figure 1), (2) the **centroid of the shell** (Figure 2b), and (3) the **centroid of the commissure** (Figure 2a). Translation to the beak positions shells onto the homologous point of initial shell growth. Translation to the centroids of the shell or its commissure incorporate more information on the shape of the shell, with centering on the commissure adding an aspect of homology by positioning shells on their growth front. Operationally, Procrustes Analysis translates landmark sets to their respective centroids before minimizing their rotational distances, overriding any prescribed translations; the three translations above are therefore implemented after the rotation step (following the functionality in *Morpho::procSym* Schlager 2017).

#### 3.2.3 Rotation

Rotation in Procrustes Analysis orients landmark coordinates to minimize their pairwise sum of squared distances. The 5 orientations discussed in the introduction were used for rotation. Because Procrustes Analysis uses Cartesian coordinates, two landmarks were placed on the mesh surface of a shell to indicate the direction of each axis as described in the subsections below (exact placement of landmarks on specimens in Figure S1).

##### Sagittal orientation

###### Sagittal axis [SX]

This axis is the pole to the commissural plane (Figure 1). It is determined as the average cross product of successive vectors that originate at the centroid of the commissure and terminate at semilandmarks on the commissure curve (visualization of fitting the commissural plane in Figure S2). The distal direction is towards the exterior surface of the shell and the proximal direction is towards the interior surface.

##### Anteroposterior orientation

###### Hinge line [HL]

The hinge line is determined by landmarks placed at the two farthest apart articulating elements of the hinge area (Figure 1). The landmarks are then designated as being anterior or posterior using the available discriminating features on the shell and can thus proxy the anteroposterior orientation. While not an ‘axis’ in the strict anatomical sense, we group the hinge line with the other anatomical axes below.

###### Oro-anal axis [OAX]

The positions of the mouth and anus or proxies thereof are used to orient the oro-anal axis and thus the anteroposterior orientation. For dimyarian taxa, anterior and posterior ends of the axis are determined by landmarks placed on the dorsal-most edge of the anterior and posterior adductor muscle scars (Figure 1a, the ‘Type 2 adductor axis’ of Bailey 2009:493 after Stanley 1970:19). For monomyarian taxa that have retained the posterior adductor muscle, the centroid of that adductor muscle scar is landmarked as the posterior end of the axis and shell features that reflect the position of the mouth are landmarked as the anterior end (e.g. the ventral notch of the anterior auricle in pectinids [Figure 1b] or the beak in ostreids, Yonge 1954). The axis is reversed in monomyarian taxa that have retained the anterior muscle (e.g. the protobranch Nucinellidae, Glover and Taylor 2013).

##### Dorsoventral orientation

###### Maximum growth axis [GX]

The origin of shell growth at the beak is the dorsal landmark on the maximum growth axis and the point on the shell commissure with the greatest linear distance to the beak is ventral landmark (Figure 1).

###### Orthogonal hinge line [oHL]

By definition, the orthogonal line to the HL (oHL) represents the dorsoventral axis, with the dorsal-most point nearest the beak.

###### Orthogonal oro-anal axis [oAX]

By definition, the orthogonal axis to the OAX (oOAX) represents the dorsoventral axis, with the dorsal-most point nearest the beak.

##### Commissure orientation

The manually placed landmarks on the shell commissure were used to fit a three-dimensional spline on which 50 equally spaced semilandmarks were placed in an anterior direction (clockwise for left valves when viewed towards the interior surface, counterclockwise for right valves). The semilandmark on the commissure curve nearest the beak landmark was selected as the initial point (Figure 2a). Semilandmarks were then slid to minimize bending energy and reduce artifactual differences in shape driven by their initial, equidistant placement (Gunz et al. 2005; Gunz and Mitteroecker 2013; Schlager 2017).

##### Standardized axis points

We observed that variance among specimens in the distances between the axis landmarks notably influenced their best-fit orientation in the Procrustes Analysis (see difference in spacing of these landmarks in Figure S1). To remove this ‘Pinocchio effect’ (*sensu* Zelditch et al. 2012:67), axis landmarks were ‘standardized’ (visualization of this process in Figure 3). The vector defined by the two axis landmarks was shifted to the centroid of the shell commissure and normalized to unit length; the standardized axis points (we explicitly avoid calling them landmarks) were then designated by the terminal points of the unit vector and its negative. Standardized axis points result in alignments that better reflect the collective impacts of axis direction, not magnitude.

**Figure 3.**
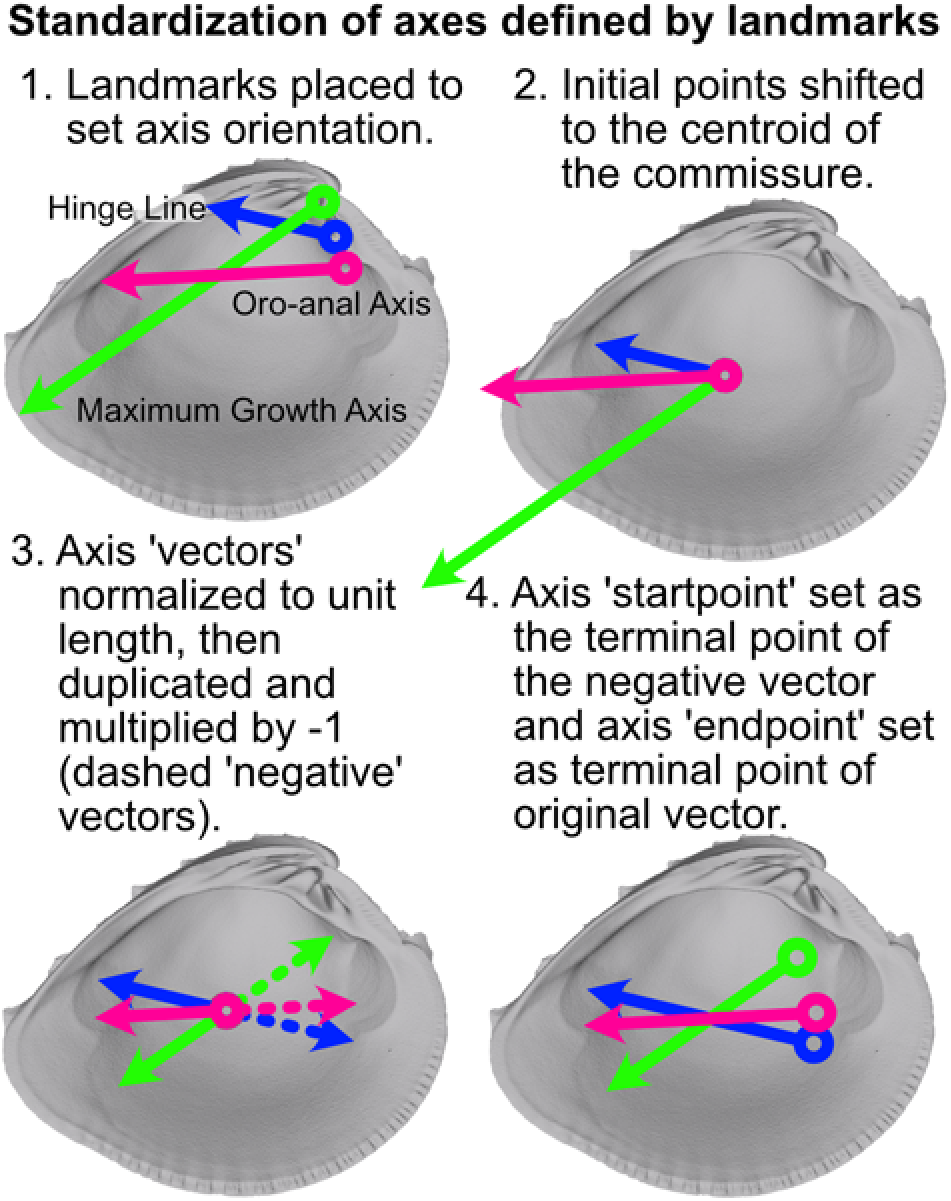
Visualization of procedure used to standardize the orientation axes defined by landmarks. Figured shell is *Chione elevata*.

### 3.3 Alignment and comparison of shape differences

Meshes and landmark sets for right valves were mirrored across their commissural plane and analyzed as operational left valves. This is a reasonable approach for equivalve taxa when analyzing general shell shape, e.g. of the interior or exterior surfaces, but homologous valves should be used for analyses that include inequivalve taxa as, by definition, their two shapes differ. Landmark sets were then scaled, rotated, and translated (in that order) under all possible parameter combinations outlined in the preceding section, totaling 45 alignment schemes. Landmark coordinate values were scaled by dividing the landmark coordinates by a specimen’s size (e.g. centroid size or volume). Scaled landmarks were then temporarily centered on the centroid of the commissure and then rotated via the respective orientation scheme using Generalized Procrustes Analysis (*Morpho:procSym*, Schlager 2017); scaling during this step was explicitly disallowed. Lastly, scaled and rotated landmarks were translated to one of the three target locations (i.e. the beak or centroids of the commissure semilandmarks or shell points).

Similarity of alignments was quantified using the metric distances between the shapes of interior shell surfaces, which were used to reduce the impact of exterior ornamentation on the differences in general shell shape. Sliding semilandmarks on the commissure and the interior surface of the shell were used to capture ‘shape.’ Initially, for the commissure, 50 equidistant semilandmarks were placed and the curve’s starting point was determined by the orientation scheme (e.g. starting at the semilandmark nearest the beak for the SX-COMM orientation, see details in Supplemental Text §2.3, Figure S5); for the interior surface, semilandmarks were placed at proportionate distances along the dorsoventral and anteroposterior axes of each orientation scheme (5% distance used here, which results in 420 semilandmarks; see details and step-by-step visualization in Supplemental Text §2.3, Figure S3, Figure S4, Figure S5). Mixing the orientations of semilandmarks and rotation axes may be useful for comparing the interaction of growth and anatomical direction with shell shape, but this approach can result in unintuitive, and perhaps unintended, shape differences among specimens. After placement of equidistant semilandmarks, those on the commissure curve were slid to minimize their thin-plate spline bending energy and then used to bound the sliding of the surface semilandmarks (Figure S5; Gunz et al. 2005; Gunz and Mitteroecker 2013; implemented via *Morpho::slider3d*, Schlager 2017). The final sliding semilandmark set consisted of 430 landmarks (50 points on the commissure plus the 380 points on the surface grid which do not lie on the commissure, i.e. the non-edge points; Figure S5). Landmark coverage analyses may be used at this point to maximize downstream statistical power (Watanabe 2018), but we relied on qualitative assessment of shape complexity and landmark coverage for the simple analyses conducted here.

For each of the 45 alignments, similarity in shell shape was calculated as the pairwise Euclidean distances of the sliding surface semilandmarks. Identical shapes have a distance of zero. Pairwise distances between shapes for each alignment scheme were normalized by their respective standard deviations, making the distances between specimens comparable across alignments. These normalized pairwise distances were then compared in three ways. First, a permutation-based multivariate analysis of variance (perMANOVA as implemented by *geomorph::procD.lm*, Adams et al. 2021) was used to model the effects of Procrustes Analysis steps on the scaled pairwise distances between specimens. Each of the 45 rows in the analyzed matrix was a unique Procrustes Analysis treatment, or alignment—i.e. a combination of scaling, rotation, and translation—and each of the 55 columns was a scaled distance between a pair of the eleven specimens. Second, this alignment matrix was scaled and centered and Principal Components Analysis was conducted to visualize the individual and joint effects treatments across alignments (i.e. Figure 4). Third, ‘hive diagrams’ (see network plots in Figure 5d) were used to compare the scaled pairwise distances of specimens among selected alignments to a reference alignment, where scaling = centroid of commissure, rotation = HL-oHL, translation = centroid of shell commissure.

**Figure 4.**
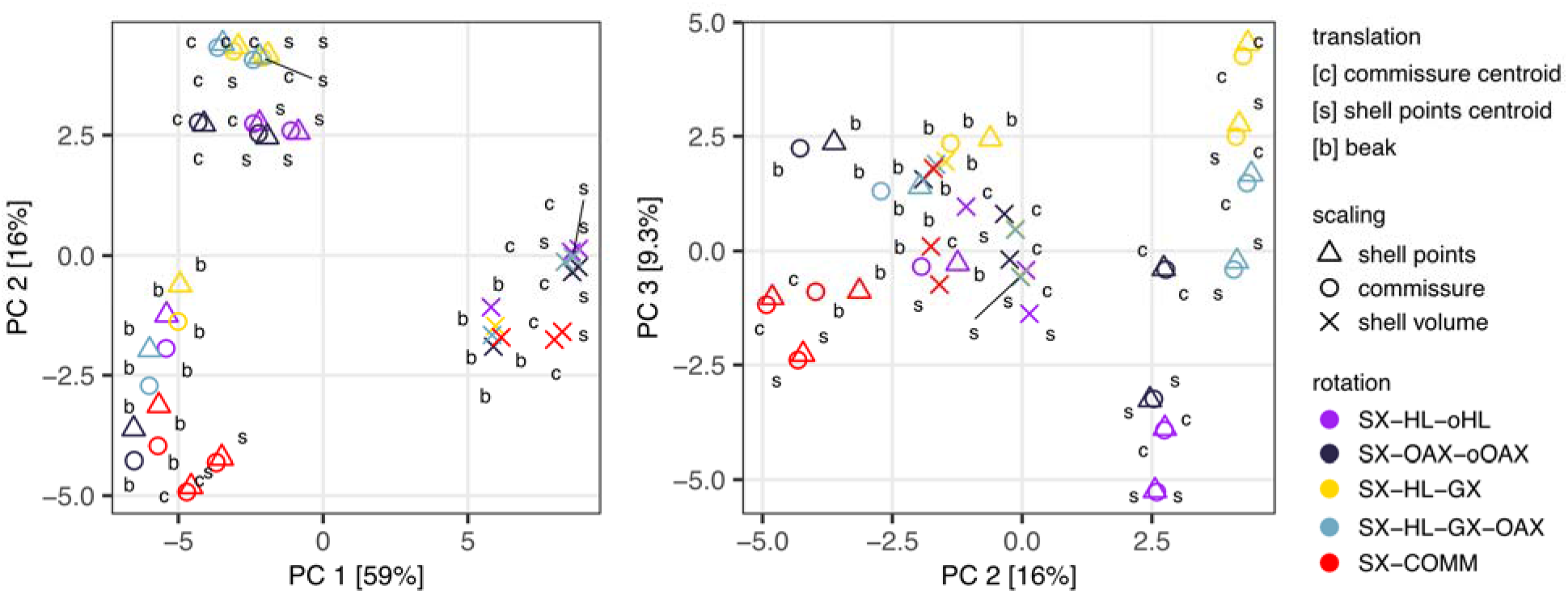
Principal Components Analysis of forty-five alignments (represented by points). Spacing of alignments along the first three axes that describe 84.3% of the total variance reflects the results of the MANOVA in Table1. Scaling by shell volume vs. the centroid size of either the shell points or the commissure separate along PC1; together, PCs 2 and 3 show the clustering of alignments with SX-COMM rotation, translation to the beak, the similarity of translation to the centroids of the commissure semilandmarks and shell points, and separation of the axis-based rotation schemes.

**Figure 5.**
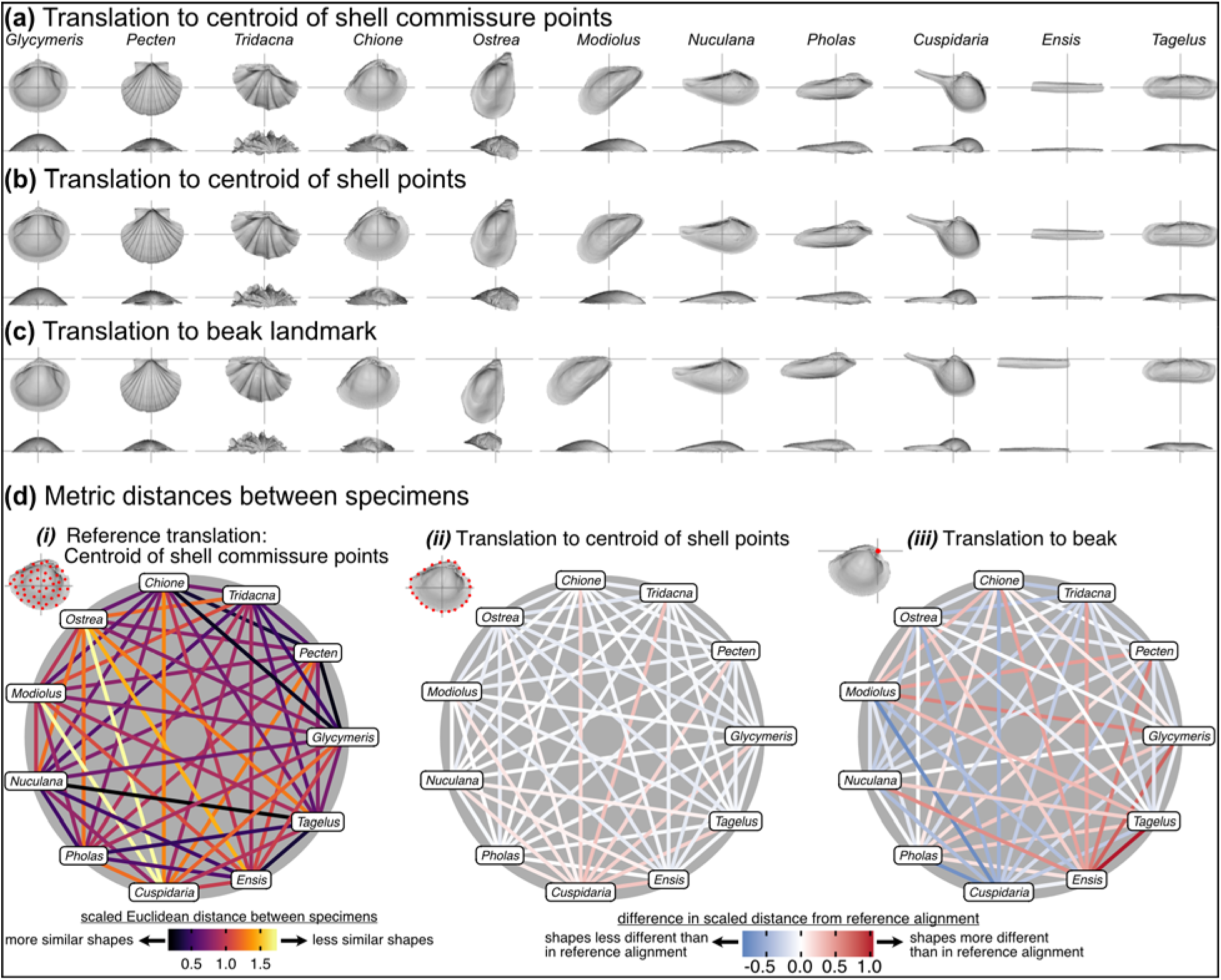
Effects of translation on differences in shell shapes. All shells are scaled to the centroid size of the shell points and rotated using the SX-HL-oHL scheme. For individual images of shells, the intersection of the gray line segments marks the origin of the Cartesian coordinate system and thus the operational ‘center’ of the shell. **(a)** Translation to the centroid of the 2000 equidistant points placed on the mesh surface of the shell. **(b)** Translation of shells to the centroid of the 50 semilandmark curve along the shell commissure. **(c)** Translation of shells to the apex of the beak landmark, the initial point of shell growth. **(d) *(i)*** The scaled pairwise Euclidean distances of semilandmarks placed on the interior surface of the shell, scaled to the centroid size of the shell points and translated to the centroid of the shell commissure. ‘Hotter’ colors indicate greater relative distances between specimens. ***(ii-iii)*** The difference in scaled distance of specimens for the specified translation from the reference treatment in panel *i*. More saturated reds indicate an increase in scaled distance relative to the reference alignment; conversely, more saturated blues indicate a decrease in distance; white indicates no difference. For example, *Ensis* and *Tagelus* become more dissimilar in interior shell shape when translated to their respective beaks than when each are translated to their centroid of the commissure.

## 4. Results and Discussion

All three Procrustes Analysis steps—translation, scaling, and rotation—significantly affect the alignments of shells (Table1, visualized as clustering of steps in Figure 4). As defined in the Methods, quantitative similarity in alignment was determined using the scaled, pairwise Euclidean distances among the sliding semilandmarks placed on the interior surface of the shell. Treatments within steps (e.g. translation to the beak vs. centroid of the shell) differentiate alignments while the effects of other parameters are held constant (*p*=0.001 for all parameters, Table1). Scaling most strongly differentiates alignments, with rotation and translation having smaller effects (see standardized effect sizes as *Z* scores in Table1). It follows that the differences between alignments are smallest among rotation and translation treatments (i.e. the smallest distances between alignments reflected as the lowest Sum of Squares, Table1, see also their clustering in Figure 4), and they increase for treatments of scaling. These metric differences among alignments are informative for understanding the impacts of individual steps in Procrustes Analysis, but visually comparing the orientations of shells is necessary to understand an alignment’s fidelity to biological homology and/or analogy.

**Table 1.** Permutation multivariate analysis of variance of the scaled pairwise distances between specimens across the 45 Procrustes Analysis alignments.

| Term | *df* | Sum of Squares | Mean Square | $R^2$ | $F$ | $Z$ | $p$ |
| --- | --- | --- | --- | --- | --- | --- | --- |
| translation | 2 | 30.6 | 15.3 | 0.11 | 13.7 | 4.4 | 0.001 |
| scaling | 2 | 185.4 | 92.7 | 0.64 | 82.7 | 6.2 | 0.001 |
| rotation | 4 | 34.7 | 8.7 | 0.12 | 7.7 | 4.8 | 0.001 |
| **Residuals** | 36 | 40.3 | 1.1 | 0.14 |  |  |  |
| **Total** | 44 | 291.0 |  |  |  |  |  |

### 4.1 Effects of translation

The choice of translation can change how differences in shell shape are interpreted. Translation to the beak, the lone homologous point across the class, allows the comparison of shell shapes conditioned on directions of growth from their origins (Figure 5c). *Ensis* can be described as being posteriorly elongated compared to *Glycymeris*, or *Pecten* as ‘taller’ than *Pholas* from the beak to the ventral margin. Still, translation to the beak can exaggerate or bias the differences in ‘pure’ shell shape. For example, *Ensis* and *Tagelus* have greater distances between their shapes when translated to the beak than to the centroid of commissure semilandmarks (red line in Figure 5d.iii). Their offset positions of the beak underlies this difference, which is interesting for analyses of growth vs. shape, but the shape of the shell, irrespective of its growth, is arguably the primary target of ecological selection (Stanley 1970, 1975, 1988; Vermeij 2002; Seilacher and Gishlick 2014). Thus, measuring the morphological similarity of shells for studies of ecomorphology, trends in disparity, or evolutionary convergence would be best conducted using translation to their respective centroids of the commissure or shell surface (Figure 5a,b); these two translations yield very similar alignments given the close proximity of their respective centroids (as shown by the pale colors linking specimens in Figure 5d.ii; but note the small offset between the two centroids for the more irregularly shaped *Cuspidaria*). In general, it is best practice to translate shells to their respective centroids of the commissure semilandmarks or shell points when morphological analyses target differences in pure shell shape. Translation to the centroid of the commissure is preferred because it incorporates homology into the alignment via correspondence of the leading edge of shell growth.

### 4.2 Effects of scaling

Scaling has a large effect on the differences between alignments (see largest Sum of Squares for the term in the ANOVA, Table1). Scaling by volume leaves particularly large residual differences in shape; the most voluminous shells are made extremely minute (*Pecten* and *Tridacna* in particular, Figure 6c) while the least voluminous shells become the largest (*Nuculana* and *Cuspidaria*). Scaling to logged shell volume does not alleviate these residual differences (results not shown), and, moreover, the aim of this scaling step is to remove the isometric relationship of size to shape, not its allometric one. The relative sizes of specimens are more similar when scaled to the centroid size of the commissure semilandmarks or the shell points (Figure 6a,b). These two sizes are tightly correlated (Figure S6) and thus produce very similar alignments (Figure 6d.ii). For comparing differences in overall shell morphology in 3D, scaling by the centroid size of shell points would best equalize the isometric differences in size among specimens, thus concentrating the remaining differences in morphology to their shapes.

**Figure 6.**
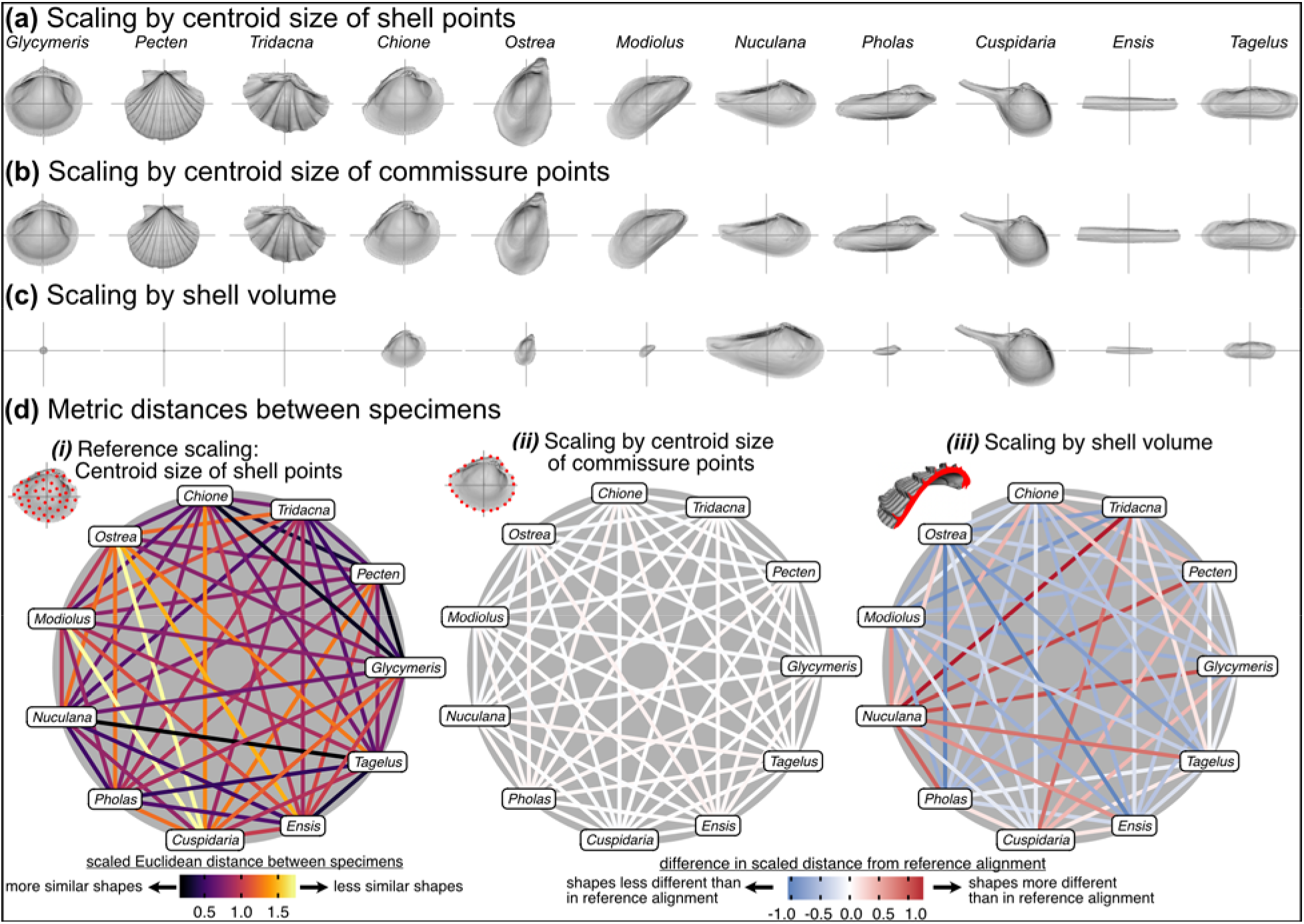
Effects of scaling on differences in shell shapes. All shells are translated centroid of the commissure semilandmarks and rotated using the SX-HL-oHL scheme. Compare differences in scaled sizes of specimens across rows, not columns. **(a)** Shells scaled by the centroid size of the 2000 equidistant points placed on the surface of the shell mesh. **(b)** Shells scaled to the centroid size of the 50 semilandmark curve along the shell commissure. **(c)** Shells scaled by the volume of shell carbonate. **(d)** As in Figure 5d but based on differences in scaling.

### 4.3 Effects of rotation

Of the three Procrustes Analysis parameters, rotation is arguably the most important factor defining the biological basis for differences in shell shape. Visually, rotation treatments can produce nearly orthogonal orientations of specimens (compare the nearly orthogonal orientation of the traditional shell length axis for *Ensis* and *Tagelus* between SX-HL-oHL and SX-COMM, Figure 7a,e; also reflected in the deep-red bar linking these two taxa in Figure 7f.v and is spacing of specimens on the first two PC axes in Figure S7). Equilateral shells are aligned similarly across rotation treatments (compare orientations of *Glycymeris*, *Pecten*, and *Tridacna*, cf. *Ostrea*, in Figure 7a-e and the less saturated lines connecting them in Figure 7e.ii-v). Differences in alignments become more pronounced among the more inequilateral shells (seen to a minor extent in *Chione* relative to *Glycymeris* and *Pecten*, but notably in *Modiolus, Pholas, Cuspidaria*, and *Ensis*) Thus, alignments of inequilateral shells tend to reflect a compromise between the often subparallel but not orthogonal orientations of their axes (most clearly seen in the changes to the orientation of *Modiolus*, *Pholas*, and *Ensis* relative to *Glycymeris* and *Pecten* in Figure 7a-f). Rotation by sliding semilandmarks on the commissure results in a similar alignment of most shells to the hinge line orientation (pale lines in Figure 7e.v), but the relative shape differences of *Modiolus, Ensis*, and *Tagelus* indicate the importance and impact of the beak position. The commissure curve begins at the point nearest the beak, which affects the orientation of the surface semilandmark grid (see Figure S5). Thus, in the SX-COMM treatment, the growth and therefore ‘shape’ of *Modiolus* and *Ensis* is more similar to the tall-shelled *Ostrea* than either are to the putative, similarly elongate *Pholas* and *Tagelus* (which themselves become more dissimilar in shape owing to the slight offset in their beak positions). Overall, rotation using the hinge axis and its orthogonal axis as the pseudo dorsoventral axis is likely the best practice for most analyses of shell shape, as discussed below.

**Figure 7.**
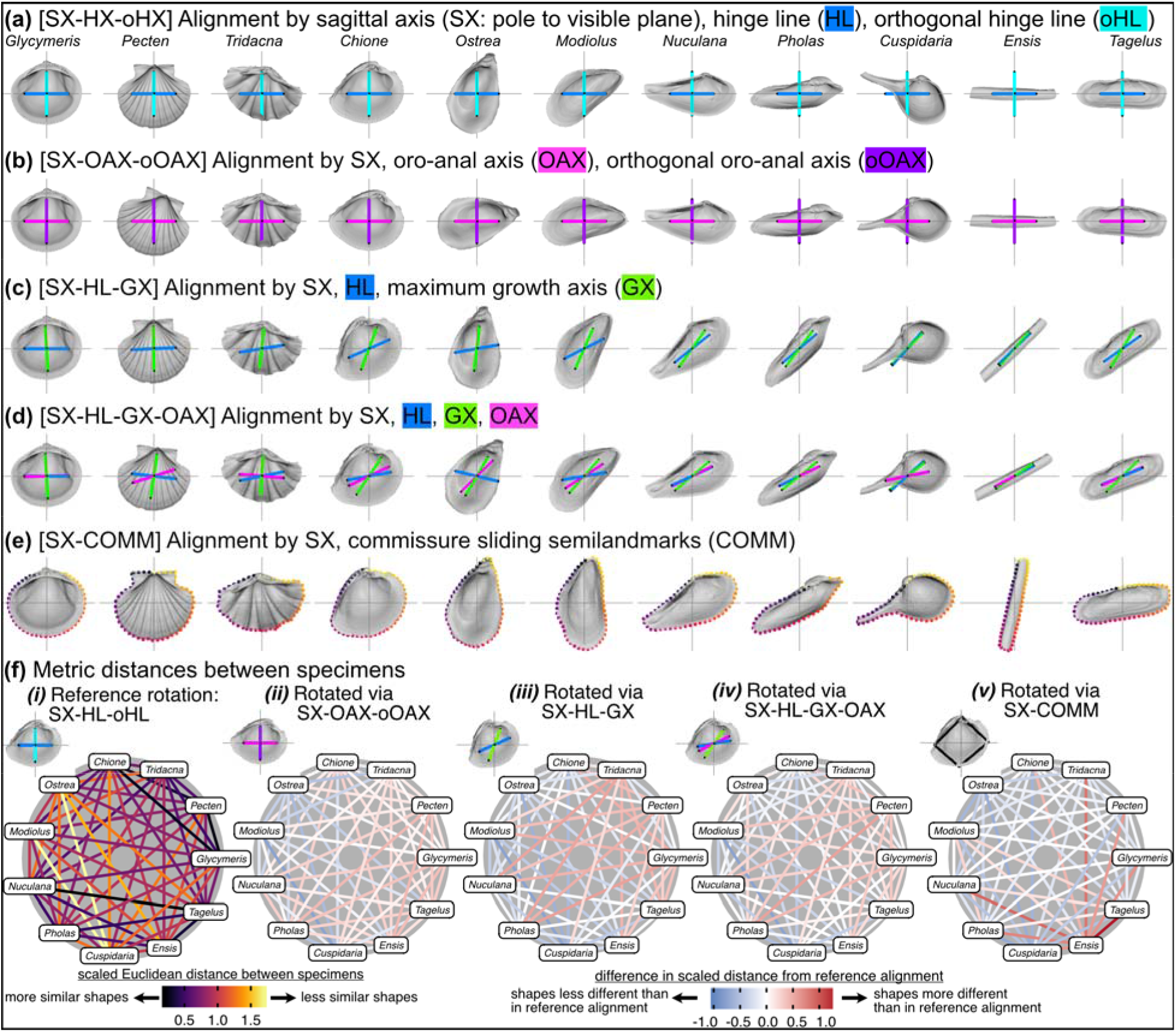
Effects of rotation on differences in shell shapes. All shells are translated centroid of the commissure semilandmarks and scaled to the centroid size of the shell points. Highlighted colors of panel titles correspond to axes plotted on shells. To facilitate relative comparisons of shell shape across columns, shells in each row were rotated such that the ‘x’ axis is parallel to the hinge line of *Glycymeris*; this is an ad-hoc, global rotation that does not change between-specimen differences in shell shape. **(a)** Shells rotated by their sagittal axis, hinge line, and orthogonal hinge line as the pseudo dorsoventral axis. **(b)** Shells rotated by their sagittal axis, oro-anal axis, and orthogonal oro-anal axis as the pseudo dorsoventral axis. **(c)** Shells rotated by their sagittal, hinge, and maximum growth axes. **(d)** Shells rotated by their sagittal, hinge, maximum growth, and oro-anal axes **(e)** Shells rotated by their sagittal axis and commissure semilandmarks. **(f)** As in Figure 5d but based on differences in rotation. See Figure S7 for a projection of shell shape differences along the first two principal components.

### 4.4 Practical alignments for bivalve shells

The axis-based approach to alignment (Figure 7a-d) is useful both for its ability to encompass broad phylogenetic analyses of shell morphology and for its ability to combine extant and fossil data, the latter known almost exclusively from shells. All shell morphologies should fit within this scheme, including those with strong lateral asymmetry (e.g. rudists, see Jablonski 2020, and those with calcified tubes or crypts (teredinids and clavagellids, Morton 1985; Savazzi 1999, each of which have identifiable valves with anatomical axes—whether to include the tubes and crypts as aspects of shell morphology is a different debate). With increasing phylogenetic proximity, the number of point-based biological homologies is likely to increase, permitting more traditional approaches to specimen alignment (Roopnarine 1995; Roopnarine et al. 2008; Márquez et al. 2010; Serb et al. 2011; Collins et al. 2013, 2020; Edie et al. 2022; Milla Carmona et al. First View). These shell-based axes and features (Figure 7a-e) are also useful for incorporating fossil taxa into analyses with extant taxa (Yonge 1954; Cox et al. 1969; Stanley 1970; Bailey 2009), but aspects of the internal anatomy remain crucial for orientation (Stasek 1963*a*), especially the designation of the anterior and posterior ends. Fortunately, in many cases the anteroposterior axis can be determined from imprints of the soft anatomy on the shell surface (e.g. the pallial sinus) or from other shell features (e.g. siphonal canals, pedal gapes). This necessarily variable and often idiosyncratic approach to defining the direction of anatomical axes may result in more digitization error than seen in traditional point-based geometric morphometrics. However, the impact of that error on analytical interpretations of shape similarity and variance will depend on the overall scale of shape disparity; for analyses of morphology across the class, the latter is likely to far exceed the former.

In biological systems with limited homology in a strict, point-based, geometric sense— and even in those with plenty of it—numerous approaches have been used to align specimens for morphological analysis. A single solution likely does not exist, and comparisons among different methods will be the most powerful approach to testing evolutionary hypotheses (see Bromham 2016 for the necessity of comparative analyses in historical science). As for most analytical frameworks, comparisons of shell shape will require explicit definition of the alignment scheme and interpretation of any differences within those boundaries. Thus, we cannot declare outright that one of these alignment schemes is logically superior, but we do recognize a practical solution that, to us, best reflects the decades of study of shell morphology: alignment via the sagittal axis, hinge line, and its orthogonal line as the pseudo dorsoventral axis (SX-HL-oHL). Shell height, length, and width have been the principal measurements for analyzing differences in shape, and long-standing, taxon-specific ‘rules’ have become entrenched in the literature and therefore influence our interpretations of the clade’s evolutionary morphology (see discussion in Cox et al. 1969:81–82 and the continued utility of these measurements in Kosnik et al. 2006). The SX-HL-oHL rotation tends to orient shells according to the defined axes of those linear measurements. Of course, precedent need not dictate the course of future work, but here, we find it reasonable to align this ‘next generation’ of shell shape analyses with the long-standing conventions in the literature, if only for comparative purposes.

## 5. Conclusions

The debate on how to align specimens is still relevant in the current era of morphometry, where comparisons of animal form are increasingly accessible in 2D, 3D, and even 4D (Boyer et al. 2016; Olsen et al. 2017; Pearson et al. 2020). But, no matter how shapes are compared, interpretations of their differences or variances should be with respect to an assumed anatomical alignment. For comparisons of disparate morphologies, particularly those that lack biological homology conducive to point-based landmarking, alignments will likely require non-standard approaches so that shape differences do not depend on geometric correspondence alone. In bivalves, anatomical axes inferred from taxon-specific features offer a class-wide approach to orientation. One set of axes in particular (HL-SX-DVX) coincides with historical approaches to their morphometry, while another offers new insight into the relationship between shell shape and shell growth (COMM-SX). Either of these solutions are valid in their own way. This philosophy of specimen alignment may be particularly relevant to other model systems in paleobiology and macroevolution that have accretionary-style growth: gastropods, brachiopods, corals, bryozoans, etc.—each with limited point-based landmarks corresponding to biological homology.

## 6. Acknowledgments

We thank FMNH and NMNH and their staff for access to the specimens used in this study.

## 8. Data Availability

Mesh models, landmark data, and code for reproducing analyses and figures available from Zenodo: 10.5281/zenodo.6326531.

## 9. Authors’ contributions

SME and KSC designed the study, SME performed analyses, all authors contributed to the writing.

## Supplemental Material

### S1. Supplemental Data

**Table S1.** Taxa used in this study and source of material.

| family | genus | species | authority | valve | museum | catalog no. | biv3d.meshid |
| --- | --- | --- | --- | --- | --- | --- | --- |
| Cardiidae | *Tridacna* | *squamosa* | (Lamarck 1819) | L | FMNHIZ | 166020 | 317 |
| Cuspidariidae | *Cuspidaria* | *rostrata* | (Spengler 1793) | L | USNM | 811161 | 1272 |
| Glycymerididae | *Glycymeris* | *glycymeris* | (Linnaeus 1758) | L | USNM | 199801 | 1683 |
| Mytilidae | *Modiolus* | *modiolus* | (Linnaeus 1758) | R | FMNHIZ | 126621 | 570 |
| Nuculanidae | *Nucula* | *pernula* | (Mueller 1779) | L | BMNH | 20180321 | 3255 |
| Ostreidae | *Ostrea* | *capsa* | J. G. F. Fischer von<br>Waldheim 1807 | R | FMNHIZ | 279417 | 138 |
| Pectinidae | *Pecten* | *maximus* | (Linnaeus 1767) | R | USNM | 25529 | 1566 |
| Pholadidae | *Pholas* | *dactylus* | Linnaeus 1758 | L | USNM | 337277 | 2380 |
| Solecurtidae | *Tagelus* | *plebeius* | (Lightfoot 1786) | L | FMNHIZ | 177579 | 769 |
| Solenidae | *Ensis* | *siliqua* | (Linnaeus 1758) | L | USNM | 27141 | 3144 |
| Veneridae | *Chione* | *elevata* | (Say 1822) | L | FMNHIZ | 176349 | 180 |

### S2. Supplemental Methods

#### S2.1. Placement of axis landmarks

**Figure S1.**
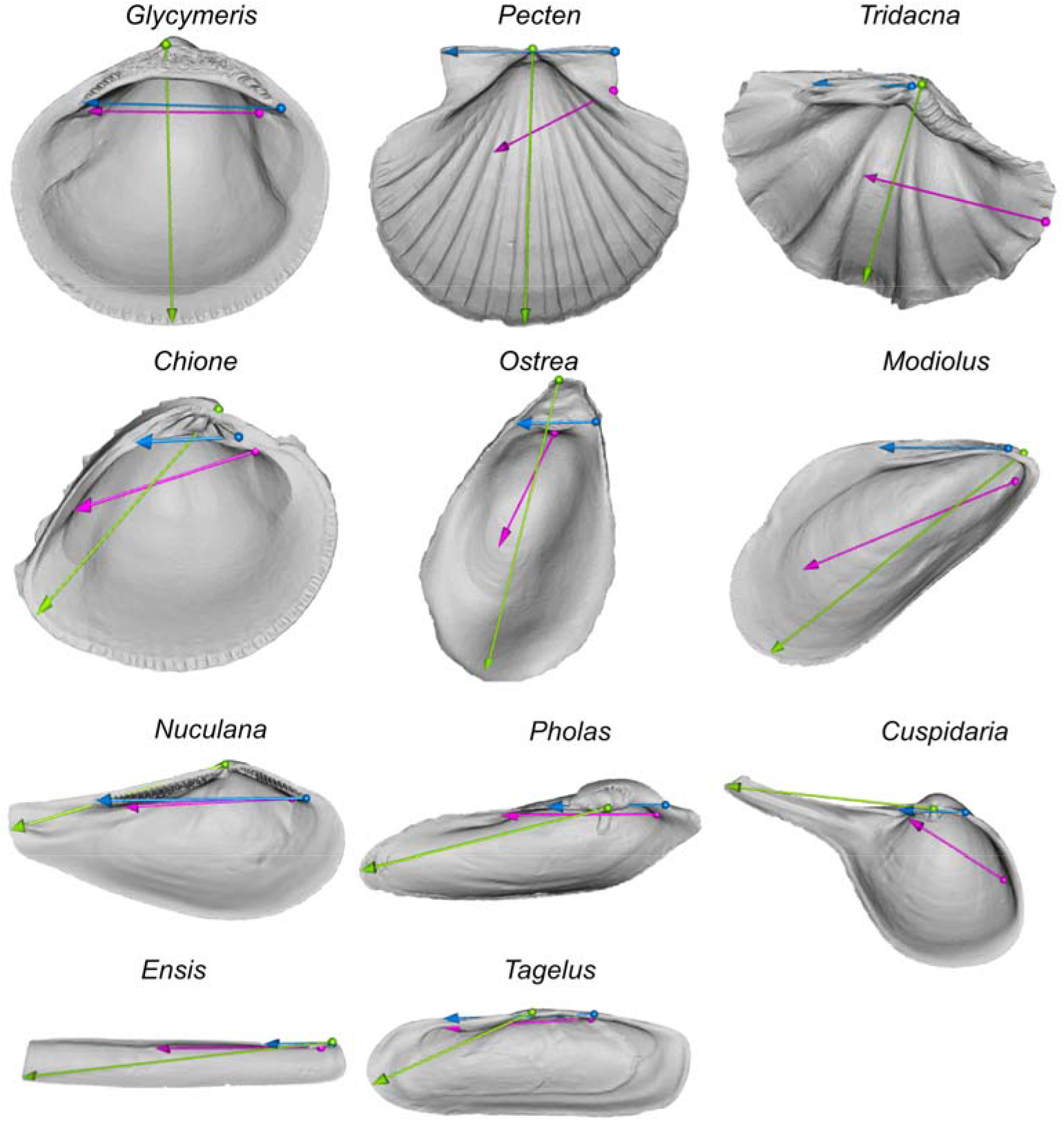
Placement of landmarks for axes (blue = hinge line; green = growth axis; magenta = oro-anal axis). Origins of axis vectors as spheres, termini as arrowheads. For the hinge line and oro-anal axis, spheres are anterior and arrowheads are posterior. For the growth axis, spheres mark the beak and arrowhead the farthest linear distance from the beak to a point on the commissure.

#### S2.2. Fitting the commissural plane

**Figure S2.**
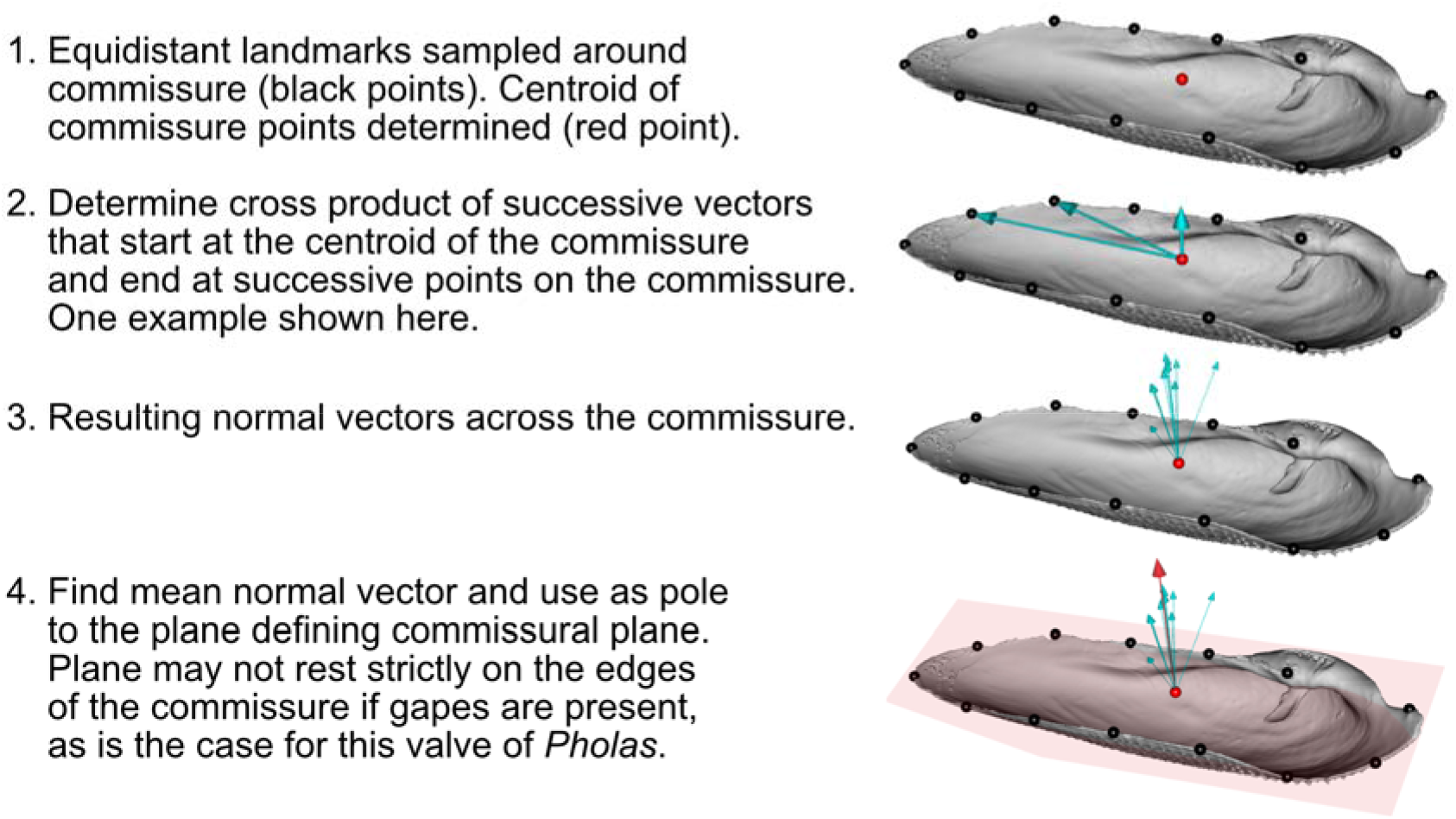
Visualization of procedure for fitting the commissural plane.

#### S2.3. Landmarking the interior surface of the shell

First, the triangular surface mesh of the shell is ‘cut’ into two pieces using the commissure curve: (1) interior (facing the commissural plane, or proximally directed on the sagittal axis) and (2) exterior (facing away from the commissural plane, or distally directed on the sagittal axis); visualization and step-by-step details in Figure S3.

Then, equidistant surface semilandmarks are placed on the ‘interior’ surface of the shell mesh as described in Figure S4 (inspired by the eigensurface method of Polly and MacLeod 2008). Note that in Figure S4-Step 5, sorting points on a flat surface best handles the ordering of points on the often topographically complex and recurved surfaces, which, in our experience, confound sorting in three-dimensions. This process is imperfect, but, again, in our experience, more reliably captures the morphology of shell surfaces compared to atlas-based approaches (Schlager 2017; Bardua et al. 2019). Bardua et al. (2019:22) state: “more accurate placement of surface points is a far more biologically sound characterization of morphology than spurious placement”—which is why we used the gridded approach in Figure S4 to place the initial semilandmarks. After placement, the equidistant semilandmarks on each individual are slid to minimize their thin-plate spline (TPS) bending energy to the mean Procrustes shape (Gunz et al. 2005; Gunz and Mitteroecker 2013; implemented via *Morpho::slider3d* Schlager 2017). The start point of the commissure curve and the orientation of the surface semilandmark grid depend on the orientation scheme:

- For the commissure orientation (SX-COMM), the initial and ‘fixed’ (i.e. non-sliding) point of the 50-point, equidistant commissure curve is the point nearest the beak (Figure S5e). The other 49 semilandmarks along the commissure curve are then slid to minimize their TPS bending energy. The surface semilandmark grid is laid down at 5% distances along an arbitrary sampling axis that spans the 13th and 38th sliding semilandmarks on the commissure curve, which generally reflect the anterior and posterior directions, respectively. The outermost grid points that intersect the commissure of the valve are removed because they will be replaced by the sliding commissure semilandmarks in the final set. The semilandmark grid is then slid to minimize its TPS bending energy, using the sliding semilandmarks on the commissure curve to constrain the sliding of the surface semilandmarks. All sliding semilandmarks are constrained to lie on the mesh surface. Thus, the final landmark set consists of 50 sliding semilandmarks along the commissure and 380 sliding semilandmarks on the interior surface of the shell, totaling 430 sliding semilandmarks.
- For the oro-anal axis orientation (SX-OAX-oOAX), the initial and ‘fixed’ point of the 50-point, equidistant commissure curve is the point that forms the smallest angle between the orthogonal oro-anal axis vector and a vector originating at the centroid of the commissure curve and terminating at a point along it (Figure S5b). The aim is to reduce the impact of the beak position on the shape of the shell, that is, to remove the effects of shell growth on comparisons on its shapes. The sampling axis for the surface semilandmarks is the oro-anal axis. The commissure curve and surface semilandarks are slid as above.
- For the orientations that include the hinge line (SX-HL-oHL, SX-HL-GX, and SX-HL-GX-OAX), the initial and ‘fixed’ point of the 50-point, equidistant commissure curve is the point that forms the smallest angle between the orthogonal hinge line vector and a vector originating at the centroid of the commissure curve and terminating at a point along it (Figure S5a,c,d). The aim is the same as for the oro-anal axis above, and the semilandmarks are slid as in the SX-COMM case above.

**Figure S3.**
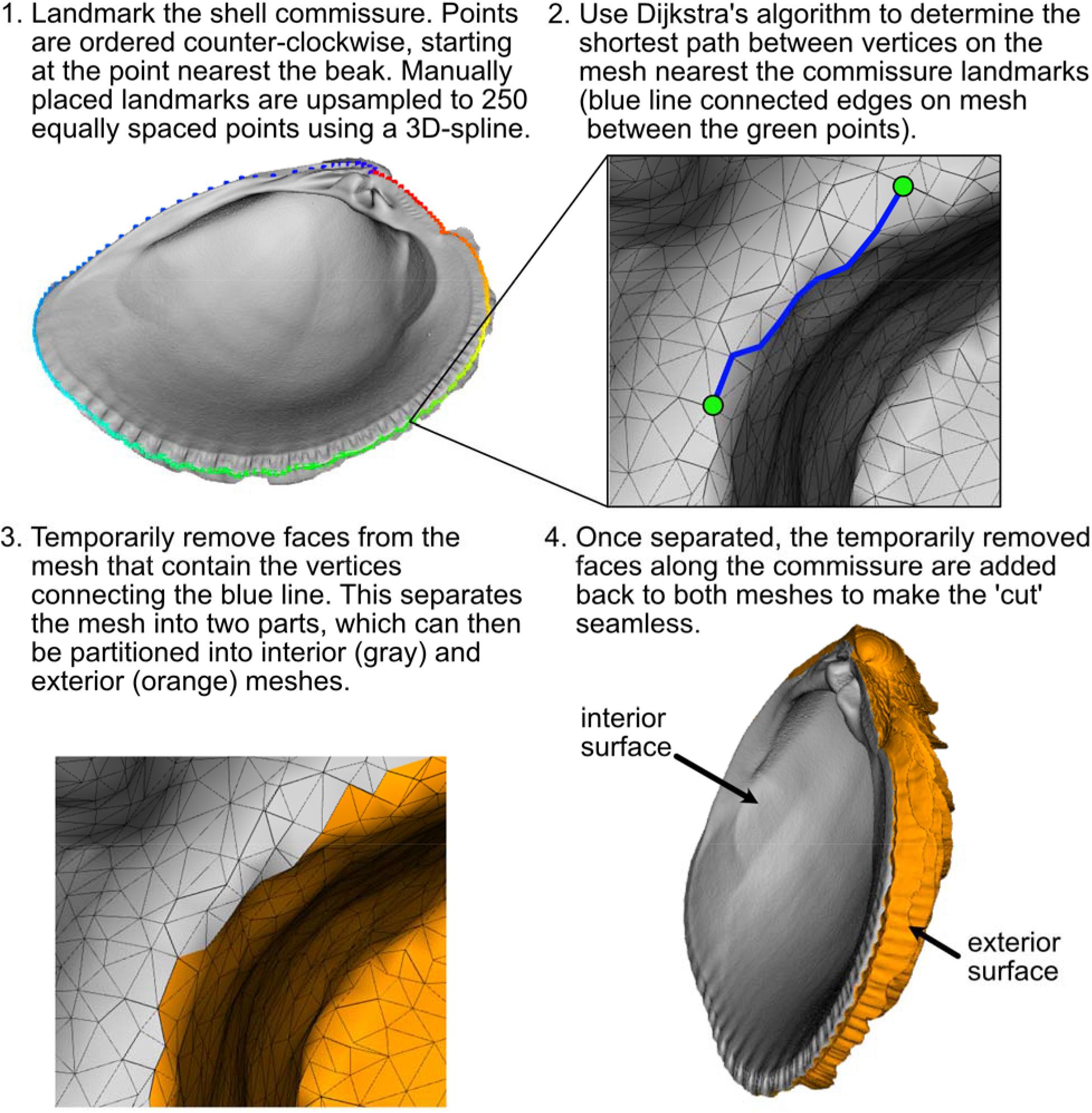
Visualization of the process for separating, or ‘cutting,’ shell meshes into interior and exterior surfaces.

**Figure S4.**
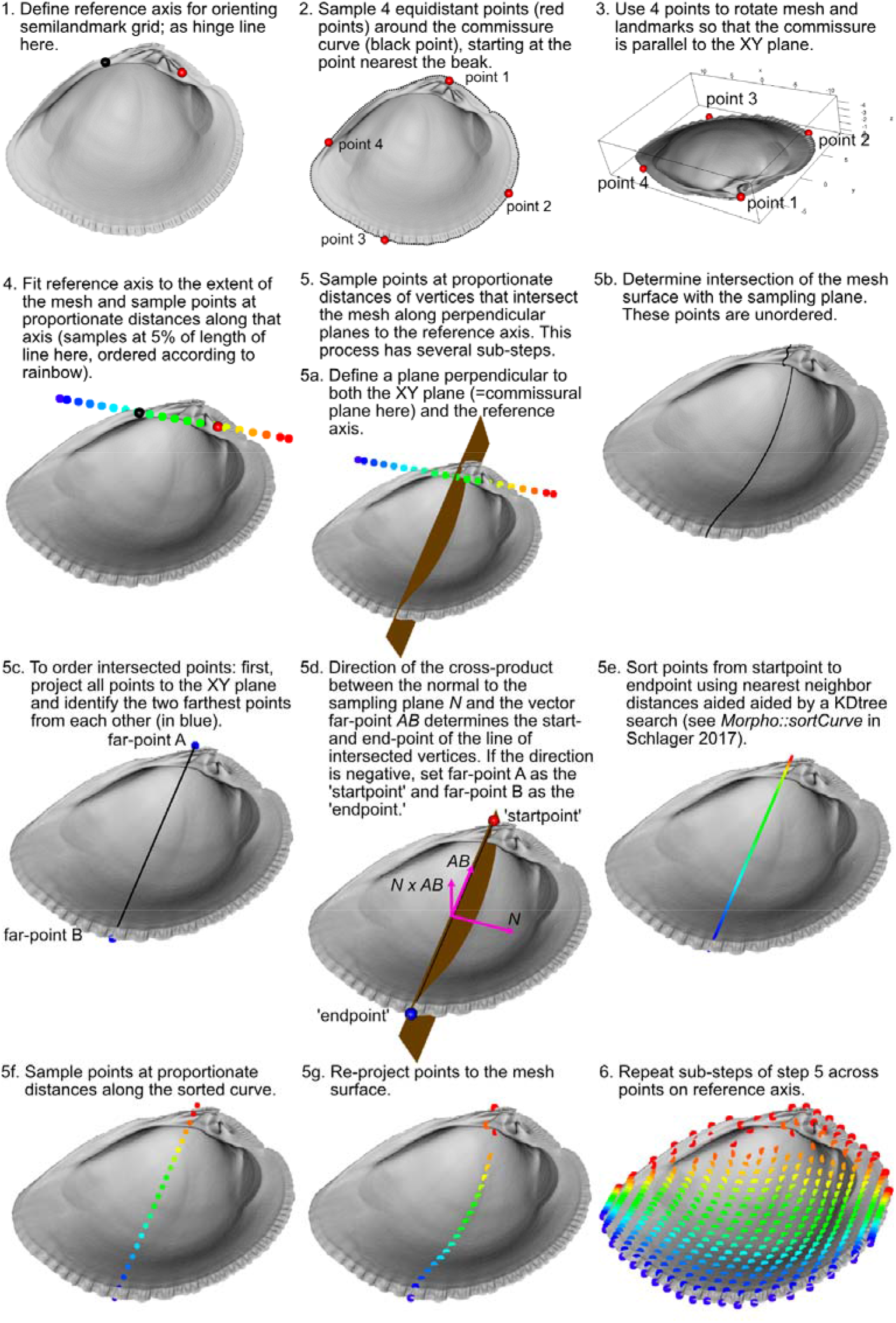
Visualization of the process for placing equidistant surface semilandmarks on the interior surface of the shell.

**Figure S5.**
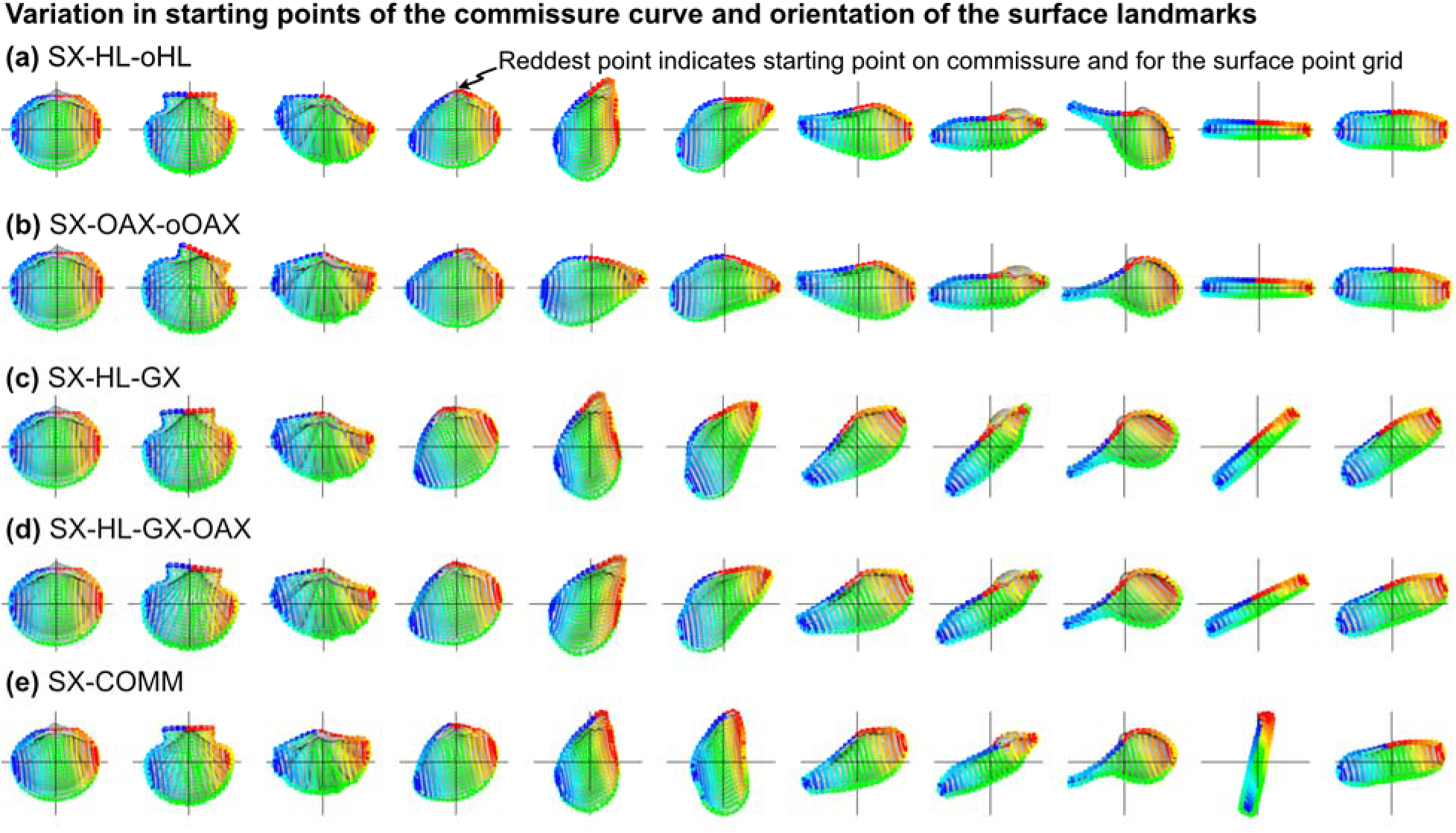
Placement of sliding semilandmarks along the commissure curve and the interior surface of the shell depending on orientation scheme. All landmark sets in this figure are scaled by the centroid size of the shell points and translated to the centroid of the shell commissure. Rainbow colored points indicate point order, with the most saturated red and blue as the respective initial and terminal points. (a) Commissure curve begins at the point that forms the smallest angle between the orthogonal hinge line vector and a vector originating at the centroid of the commissure curve and terminating at a point along it. Surface semilandmarks are oriented orthogonal to the hinge line. (b) Commissure curve begins at the point that forms the smallest angle between the orthogonal oro-anal axis vector and a vector originating at the centroid of the commissure curve and terminating at a point along it. Surface semilandmarks are oriented orthogonal to the oro-anal axis. (c) Commissure curve and surface landmarks oriented as in panel a. (d) Commissure curve and surface landmarks oriented as in panel a. (e) Commissure curve begins as the point nearest the beak. Surface semilandmarks are oriented orthogonal to the line connecting the 13th and 38th sliding semilandmarks on the commissure curve, which generally reflect the anterior and posterior directions.

### S3. Supplemental Results

**Figure S6.**
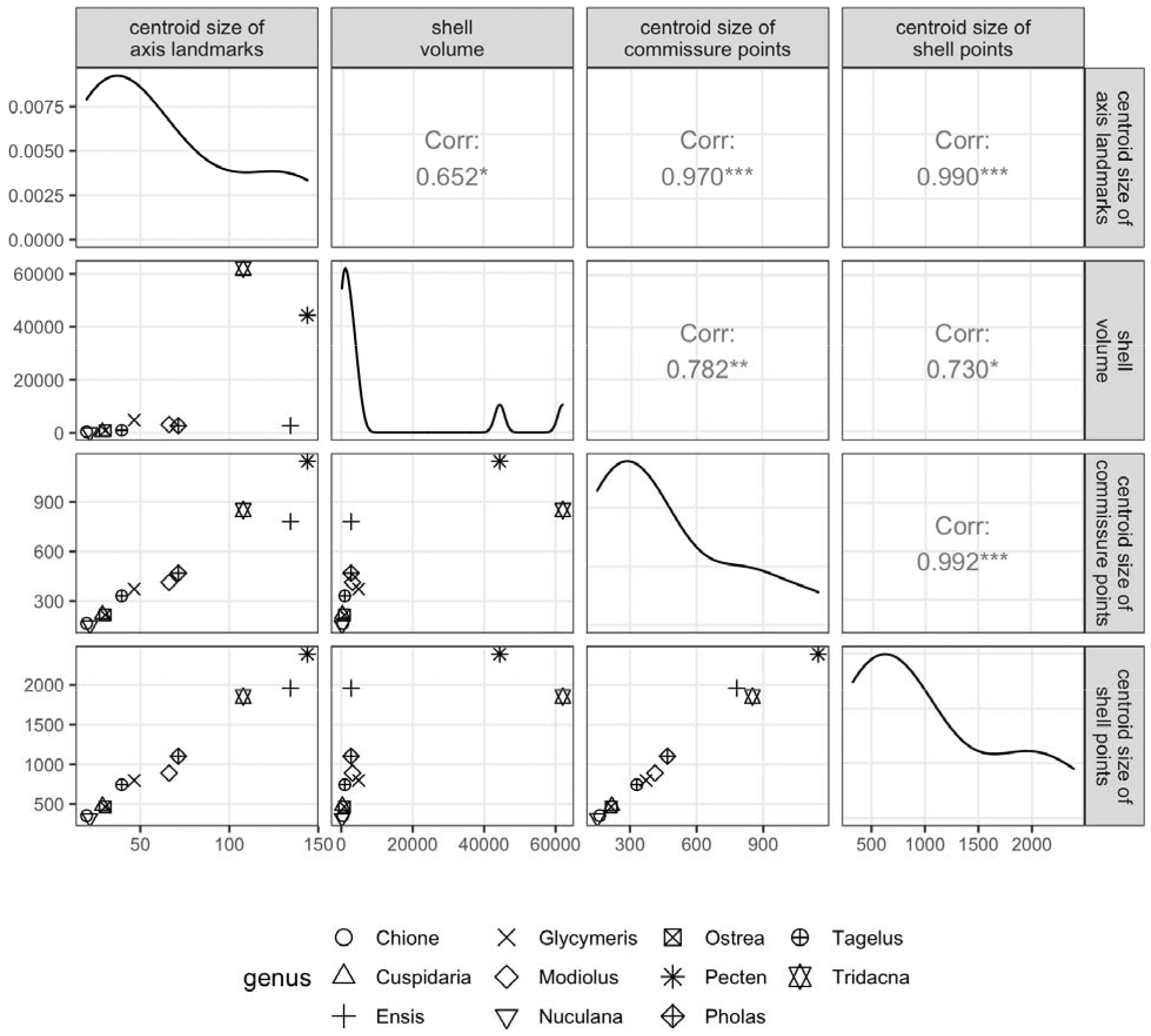
Correlations of size measures. Lower left triangle of the plot matrix shows the pairwise, bivariate relationships of size measures among analyzed specimens. Diagonal of the plot matrix shows density function for each size measure. Upper right triangle of the plot matrix shows results of Pearson correlation tests, with asterisks denoting significance at the following *p* levels: * = 0.05, ** = 0.01, and *** = 0.001.

**Figure S7.**
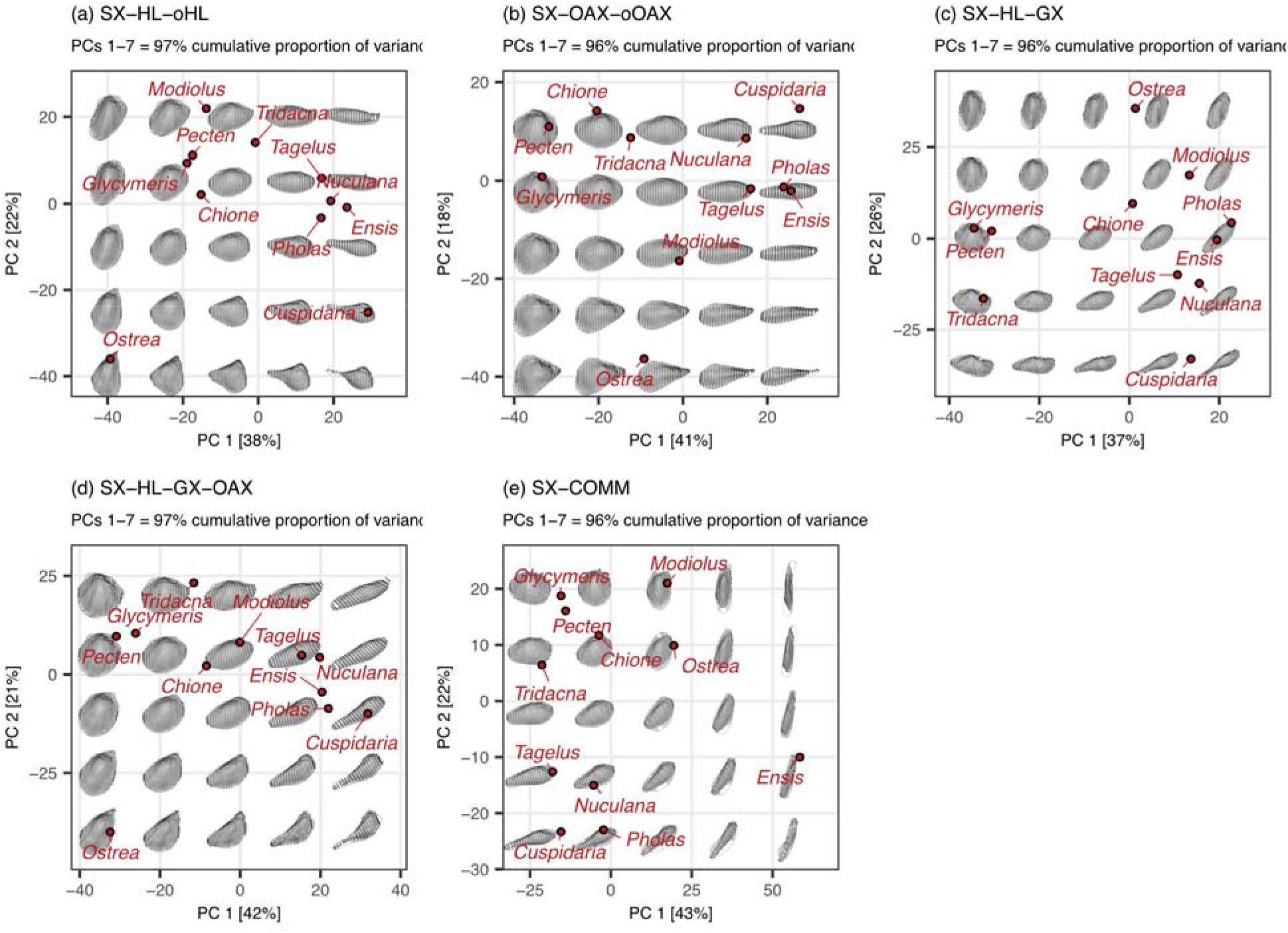
Principal components analysis of the aligned sliding semilandmarks on the commissure and interior surface of the shell. All landmark sets in this figure are scaled by the centroid size of the shell points and translated to the centroid of the shell commissure. Panels a-e give the positions of specimens on the first two principal components (PCs; percentages in brackets on each axis give the proportion of total variance explained by that axis). Images of shells are projections of the shapes at their given locations in the PC1-PC2 space. Holes in the mesh surfaces are artifacts of the meshing algorithm; the black points are the true underlying data.

